# Driving Hierarchical Collagen Fiber Formation for Functional Tendon, Ligament, and Meniscus Replacement

**DOI:** 10.1101/2020.08.07.241646

**Authors:** Jennifer L. Puetzer, Tianchi Ma, Ignacio Sallent, Amy Gelmi, Molly M. Stevens

## Abstract

Hierarchical collagen fibers are the primary source of strength in musculoskeletal tendons, ligaments, and menisci. It has remained a challenge to develop these large fibers in engineered replacements or *in vivo* after injury. The objective of this study was to investigate the ability of restrained cell-seeded high density collagen gels to drive hierarchical fiber formation for multiple musculoskeletal tissues. We found boundary conditions applied to high density collagen gels were capable of driving tenocytes, ligament fibroblasts, and meniscal fibrochondrocytes to develop native-sized hierarchical collagen fibers 20-40 µm in diameter. The collagen fibers organize similar to native collagen with native fibril banding patterns and hierarchical fiber bundles 50-350 µm in diameter by 6 weeks. Mirroring fiber organization, tensile properties of restrained samples improved significantly with time, reaching ∼1 MPa. Additionally, tendon, ligament, and meniscal cells produced significantly different sized fibers, different degrees of crimp, and different GAG concentrations, which corresponded with respective native tissue. To our knowledge, these are some of the largest, most organized fibers produced to date *in vitro*. Further, cells produced tissue specific hierarchical fibers, suggesting this system is a promising tool to better understand cellular regulation of fiber formation to better stimulate it *in vivo* after injury.

## 1. Introduction

Collagen fibers are the primary source of strength in connective tissues throughout the body, particularly musculoskeletal tissues such as tendons, ligaments, and menisci. These tissues are characterized by large hierarchically organized type I collagen fibers which assemble from tropocollagen molecules into fibrils (10 - 300 nm, up to 1 µm diameter), fibers (>10 µm diameter) and fascicles (100s µm to mm diameter, **Figure 1**) [1–3]. In tendons and ligaments these hierarchical fibers run the length of the tissue, increasing in size to form the majority of the tissue and provide the strength necessary to translate loads from muscle-to-bone and bone-to-bone, respectively [2,4]. In menisci, these fibers run circumferentially around the outer two-thirds of the tissue, providing the mechanical strength necessary to withstand and distribute compressive loads across the knee [5,6].

**Figure 1:**
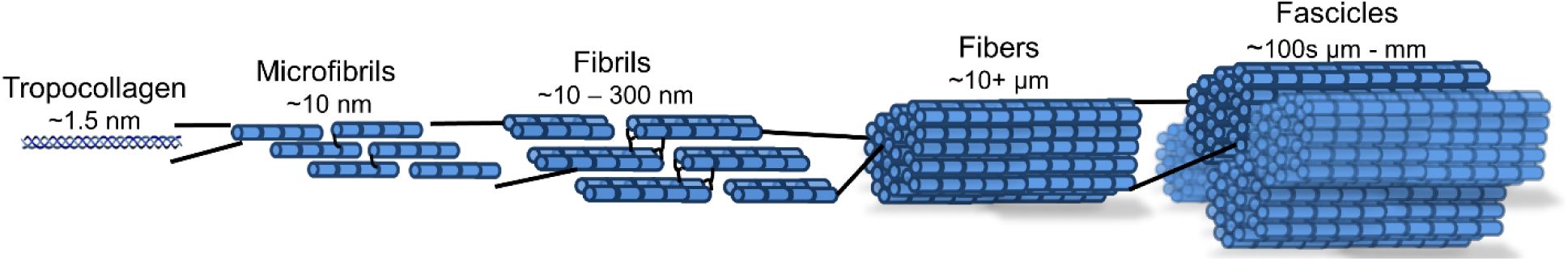
Hierarchical collagen organization with relative diameters of sub-level organization.

The hierarchical collagen fiber organization of musculoskeletal tissues is crucial to the mechanical integrity and function of the tissues. Injuries to these tissues disrupt the collagen organization resulting in loss of function, pain, and decreased mobility [4,6,7]. Due to limited blood supply and demanding load environments, these tissues are characterized by little to no healing. Ultimately, injuries to these tissues result in over 1.4 million surgeries a year in the US (based on 1,000,000 meniscus, 130,000 ACL, 275,000 rotator cuff repairs per year) [4,6–8]. Current treatments for severe tears primarily consist of autograft or allograft transplants, which have limited availability, risk of immune response, and donor site morbidity [4,6–8]. Engineered replacements are a promising alternative [4,6,7]; however, these replacements often fail to produce sufficient collagen, let alone the organized collagen fibers essential to long term mechanical success and thus often do not translate to the clinic.

Efforts to produce aligned hierarchical collagen fibers for dense connective tissues, such as tendons, ligaments, and menisci, have largely concentrated on the use of mechanical boundary conditions to guide cells to produce aligned fibrils [9–19]. These systems primarily consist of cell-seeded low density collagen or fibrin gels (1-5 mg/mL) restrained to harness cellular contraction and guide the alignment of collagen fibrils. These culture systems are well established to produce organized collagen fibrils resembling embryonic tissue (<1 µm in diameter) [11,15–19]; however due to the low density of material they undergo significant contraction, resulting in a lack of therapeutic applications and early rupture. Additionally, the fibrils do not mature into larger fibers, often resulting in inferior mechanical properties compared to native tissue. It has remained a challenge to transition from embryonic-like fibrils to the larger fibers and fascicles that dominate mature tissue.

Recently, native sized 30-40 µm diameter collagen fibers were developed in a whole meniscus system using mechanical boundary constraints and high density collagen gels (10 – 20 mg/ml) [20–22]. These are some of the largest fibers developed to date in engineered tissues, however the mechanical properties were lacking in comparison to native tissue, suggesting a need for further maturation. It has been shown that as the concentration of collagen gels increase, cellular contraction of the gel decreases and mechanical properties increase [17,20,23,24]. These attributes of high density collagen gels may provide the time and mechanical signals needed to support the development of larger hierarchical fibers, which have not been observed in low density gels. We hypothesized that high-density collagen gels could not only support further fiber development with meniscal fibrochondrocytes, but also support fiber formation with tenocytes and ligament fibroblasts. The objective of this study was to create a simplified culture system with high density collagen gels to drive cellular hierarchical fiber formation in multiple musculoskeletal tissues, and to investigate fiber maturation at the fibril, fiber, and tissue level in an effort to create functional musculoskeletal tissue with native organization and strength.

## 2. Material and Methods

### 2.1 Construct Fabrication and Culture

#### 2.1.1 Cell Isolation

Bovine tenocytes, ligament fibroblasts, and meniscal fibrochondrocytes were isolated as previously described [20,21,23]. Briefly, 2-6 week old calf legs were purchased from a local abattoir within 48 hours of slaughter to obtain deep flexor tendons of the metacarpal joint, the anterior cruciate ligament (ACL), and menisci of the knee. All tissues were aseptically isolated from the joint, diced, and digested overnight. Tenocytes were isolated from deep flexor tendons, and ligament fibroblasts were isolated from the anterior cruciate ligament (ACL) with 0.2% w/v collagenase digestion overnight [11]. Meniscal fibrochondrocytes were isolated from the entire meniscus with 0.3% w/v collagenase digestion overnight [20,21,23]. The next day cells were filtered, washed, counted and immediately seeded into high density collagen gels to avoid effects of passaging. Each cell type had 4 separate isolations (12 total) divided equally between clamped and unclamped cultures, with cells pooled from 3-4 cows each time to limit donor variation and to obtain enough cells (N = 40 cows total, 12 for flexor tendons, 16 for ACL, and 12 for meniscus).

#### 2.1.2 High Density Collagen Construct Fabrication

Cell seeded high density collagen gels were fabricated as previously described [20,21,23]. Briefly, type I collagen was extracted from purchased post-mortem adult mixed gender Sprague-Dawley rat tails (Seralab, West Sussex, UK) and reconstituted at 30 mg/mL in 0.1% v/v acetic acid [23,24]. To fabricate constructs the stock 30 mg/ml collagen solution was mixed with appropriate volumes of 10x phosphate buffer saline (PBS), 1 N NaOH, and 1x PBS to begin gelation and raise pH and osmolarity, as previously described [20,21,24]. This collagen solution was immediately mixed with cells, injected between glass sheets 1.5 mm apart, and gelled at 37 °C for 1 hour to obtain 20 mg/mL collagen constructs at 5×10^6^ cells/mL. Rectangles (8 × 30 mm) were cut from sheet gels and cultured clamped or unclamped for up to 6 weeks (**Figure 2A**). Flexor tenocytes, ACL ligament fibroblasts, and meniscal fibrochondrocytes were all cultured in media composed of DMEM, 10% v/v fetal bovine serum, 1% v/v antibiotic antimycotic, 50 µg/mL ascorbate, and 0.8 mM L-proline, with meniscal fibrochondrocytes constructs receiving an additional 0.1 mM non-essential amino acids. Media was changed every 2-3 days.

**Figure 2:**
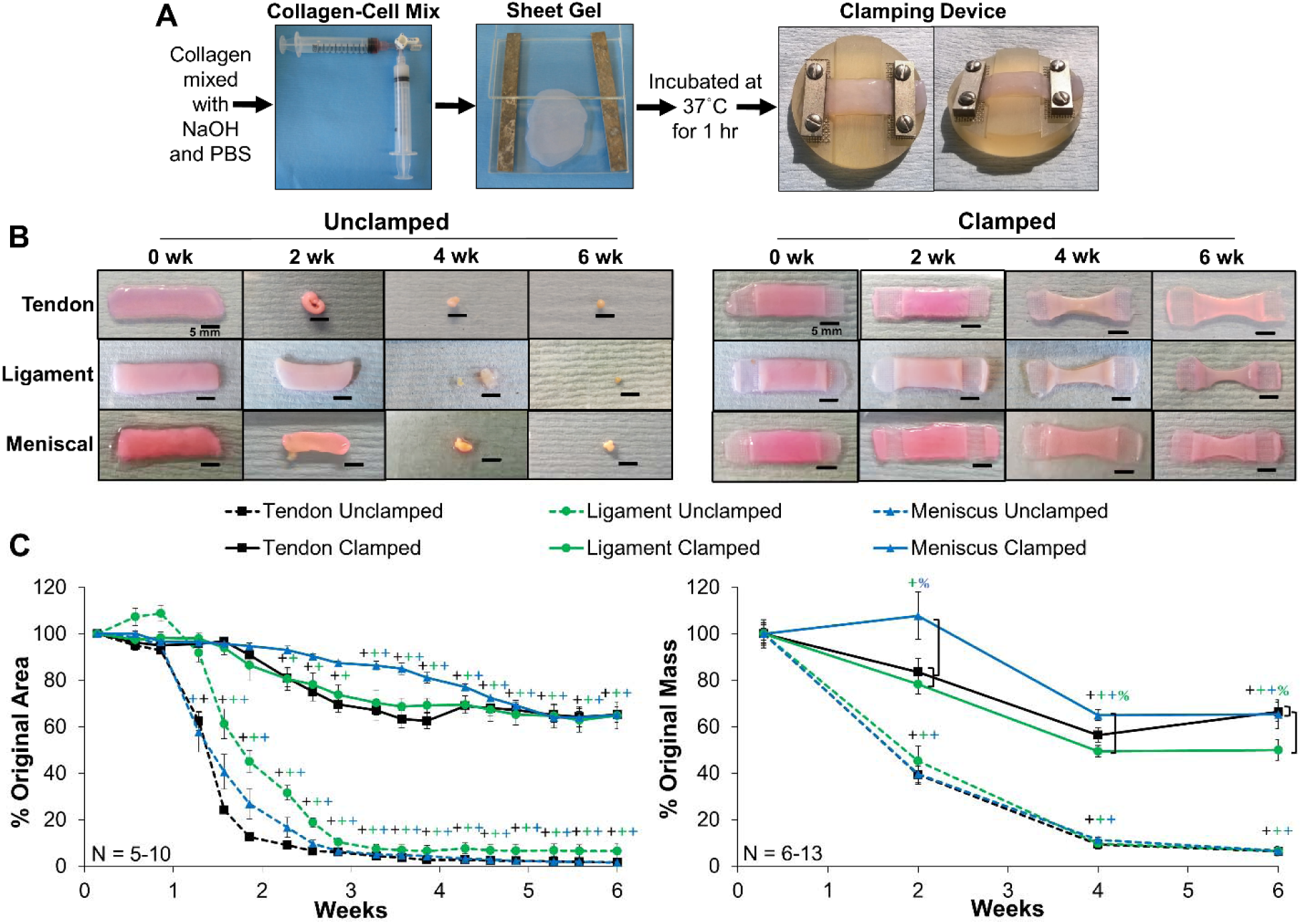
A) Cell-seeded collagen sheet gel formation and application of mechanical constraints of clamped constructs. B) Photographs of unclamped and clamped samples throughout culture and C) percentage of original area and mass of constructs. Unclamped samples significantly contracted from day 9 on, while clamped samples maintained their size through ∼2-3 weeks. Scale bar = 5 mm, Data shown as mean ± S.E.M., Significance compared to ^+^0 week and ^%^bracket group (*p* < 0.05).

#### 2.1.3 Application of Boundary Conditions and Culture

Boundary conditions were applied by a custom clamping system designed to fit within a 6 well culture dish (**Figure 2A**). It consists of a polysulphone base with 1 mm indentions on each side that allows for secure anchoring of the gels. Stainless steel clamps were screwed down over the ends of the gel to secure them throughout culture, with an additional stainless steel mesh to prevent slippage. The clamps provided boundary constraints, which limited cellular contraction and guided alignment of collagen fibrils. Clamped constructs were clamped on day 1 and remained clamped the duration of culture.

Constructs were cultured for up to 6 weeks clamped or unclamped. At 0, 2, 4, and 6 weeks constructs were removed from culture, weighted and sectioned into pieces that were fixed or frozen for analysis of collagen organization, composition, and mechanical properties. Zero week constructs were harvested on day 2 to allow for 24 hours of clamped culture conditions. Serial photographs taken throughout culture were analyzed with FIJI (Image J, NIH) to calculate area of constructs, normalized to area on day 1. The final weights of the constructs were normalized to the average weight at 0 weeks to determine change in mass with time. A variable number of constructs (N = 6 -13) were obtained for each cell type and time point due to significant contraction with time in culture, a large number of analysis techniques, and an unpredictable number of cells from each isolation. Additionally, 4 native samples were obtained for each tissue to serve as controls for collagen organization and tissue composition. N noted in figures represents number of engineered samples.

### 2.2 Collagen Organization Analysis

Full length sections were cut from each construct and fixed in 4% v/v formaldehyde, stored in 70% ethanol, and imaged with confocal reflectance to evaluate collagen fiber organization. After evaluation with confocal reflectance, a subset of 0 and 6 week constructs and native tissue were analyzed with polarized picrosirius red imagining (N = 4) and atomic force microscopy (AFM, N = 3) to evaluate fascicle and fibril level organization, respectively.

#### 2.2.1 Confocal Analysis

Confocal reflectance analysis was performed as previously described [20,21,23]. Imaging was performed with a Leica SP5 inverted confocal microscope equipped with an Argon 488 nm laser, and a HCX PL APO CS 40 ⨯ 1.25-0.5 oil objective. Noise was reduced by collecting images at 1024 ⨯ 1024 pixels with a line average of 4. Confocal reflectance was performed in conjunction with fluorescence by splitting a 488 nm laser to visualize collagen organization and cells, respectively. Confocal reflectance microscopy was performed by collecting backscatter light reflected by collagen fibers through a 40 µm pinhole at 449 - 471 nm. Auto-fluorescence of the cells was captured at 500 - 571 nm. Three dimensional images were obtained by compiling z-stacks with < 0.8 µm step size and 15 – 30 µm depth. Z-stacks were visualized with FIJI plugin 3D viewer (NIH).

Representative images were taken across the entire full length section of the construct, with sections taken from both the outside edge and center of constructs to ensure analysis of all areas. Images were analyzed with a custom Fast Fourier transform (FFT) based MATLAB [20,21] code to determine the degree of alignment (a value of 1 = unorganized, 4.5 = completely aligned) and mean fiber diameter. Three to sixteen images per sample were analyzed (depending on size of sample) and averaged, with 6-12 scaffolds analyzed per time point.

#### 2.2.2 Polarized Picrosirius Red Imaging

Fixed construct sections were embedded into paraffin, sectioned, and stained with picrosirus red. Images were taken with polarized light at 10x to observe collagen fascicle organization. Imaging was performed with a Nikon Ts2R-FL microscope equipped with a polarizer and a NIS color camera.

#### 2.2.3 Atomic Force Microscopy Analysis

Flat samples for AFM analysis were obtained as previously described [25]. Briefly, fixed samples were glued onto glass slides with a small quantity of silicone glue. They were then immersed in PBS and frozen on a cryo-sectioning block. Using a cryostat (Bright Instruments Ltd.) the tissue was cut at an angle of 2.5° and 15 µm sections to obtain a flat surface. Once a flat tissue cross section was obtained, the samples were stored in PBS and dried with nitrogen prior to AFM imaging. AFM scanning was performed with a Keysight 5500 AFM, using a qp-BioAC probe (Nanosensors TM, Switzerland) with a nominal spring constant of 0.06-0.18 N/m in contact mode or HQ NSC probe (MikroMasch, Bulgaria) with a nominal spring constant of 45 N/m in AC mode. Fibril diameters and banding (d-period length) were analyzed with Gwyddion (http://gwyddion.net). Fibril diameters were measured on 5 random fibrils per sample, with 3 cross-sectional measurements averaged along each fibril (N = 2-3 for 6 week clamped samples and N = 1 for the native tissue). Fibril d-period length (banding) was measured on 5-6 random fibrils per sample, averaging the length of the d-period along each individual fibril (on average 3-5 d-periods for 6 week clamped, 5-8 for native per fibril). Average fibril diameters and d-period length per fibril were pooled between samples in each group for general analysis of fibril characteristics (n = 10-15 for engineered, n = 5-6 for native).

### 2.3 Biochemical Composition

Portions from each engineered construct and native tissue were weighed wet (WW), frozen, lyophilized, weighed dry (DW), and digested in papain for 16 hours for biochemical analysis. DNA, collagen, and glycosaminoglycan (GAG) accumulation were determined using Quant-iT PicoGreen dsDNA assay kit (Invitrogen), a hydroxyproline (hypro) assay [26], and a modified 1,9-dimethylmethylene blue (DMMB) assay at pH 1.5 [27], respectively. Biochemical properties were normalized to WW and DW of the sample. DNA, GAG, and hydroxyproline content reported per construct were determined by multiplying the WW normalization by the total wet weight of the construct.

### 2.4 Mechanical Properties

Full length sections were cut from each construct and frozen to determine tensile properties as previously described [20,21]. All mechanical tests were performed using an EnduraTEC ElectroForce 3200 System (Bose) outfitted with a 250 g load cell. Full length strips where thawed in PBS with EDTA-free protease inhibitors, measured, and secured in clamps with a small amount of cyanoacrylate glue. Samples were tested at a strain rate of 0.75%/s, assuming quasi-static load and ensuring failure occurred between the grips. The reported tensile modulus is the slope of the linear region of the stress-strain curve as determined by a linear regression fit ensuring an r^2^ > 0.999. The end of the toe region and yielding point were determined as the point where the linear regression dropped to r^2^ < 0.999. The ultimate tensile strength (UTS) and strain at failure are the maximum stress point. Unclamped constructs could not be tested past 2 weeks due to significant contraction.

### 2.5 Statistics

SPSS was used to confirm normality of data within each group and detect outliers using Shapiro-Wilk tests and Q-Q plots. All image analyses, biochemical, and tensile modulus data were analyzed by 2 and 3-way ANOVA using Tukey’s t-test for post hoc analysis (SigmaPlot 14). Engineered 6 week clamped mechanical stress and strain data were analyzed by 1-way ANOVA with Tukey’s t-test for post hoc analysis (SigmaPlot 14). Biochemical and collagen fiber diameter correlations were analyzed by Pearson’s correlation. For all tests, *p* < 0.05 was considered the threshold for statistical significance. All data are expressed as mean ± standard error (S.E.M.).

## 3. Results

### 3.1 Construct Appearance and Shape Fidelity

Gross inspection revealed clamped scaffolds maintained their size through approximately 2-3 weeks, leveling off at 60-70% their original size and weight in the final 2 weeks. Unclamped samples significantly contracted from day 9 on, leveling off at 6-7% their original weight (**Figure 2B & C**). Interestingly, clamped tendon and ligament samples significantly reduced their size and weight by 2 weeks, while clamped menisci maintained size and weight through 3 weeks, suggesting initial differences in contraction strength and activity between cell types. All clamped samples plateau in size and weight in the final two weeks suggesting a transition from collagen reorganization to production.

### 3.2 Induced Hierarchical Collagen Organization

Confocal reflectance, polarized picrosirius red imaging, and AFM analysis demonstrated that tendon, ligament, and meniscus clamped constructs developed hierarchical collagen organizations similar to native tissue by 6 weeks of culture (**Figure 3-5**). Confocal reflectance revealed that all unclamped scaffolds remained unorganized throughout culture, while clamped constructs (tendon, ligament, and meniscus) developed aligned collagen fibrils by 2 weeks which developed into fibers (20-40 µm in diameters) similar to their respective native tissue by 6 weeks (**Figure 3A**). Further, 3D reconstructions of 6 week clamped samples revealed clearly defined collagen fibers (**Figure 3B**). Image analysis revealed all clamped constructs significantly improved collagen alignment by 2 weeks to match respective native alignment and significantly increased mean collagen fiber diameter with time in culture to match native fibers by 6 weeks. Interestingly, by 6 weeks clamped tendon, ligament, and meniscus constructs had developed significantly different collagen fiber diameters (22 ± 3 µm, 30 ± 4.4 µm, 37 ± 4.6 µm in diameter, respectively; *p <* 0.001), which matched their respective native tissues (**Figure 3C**).

**Figure 3:**
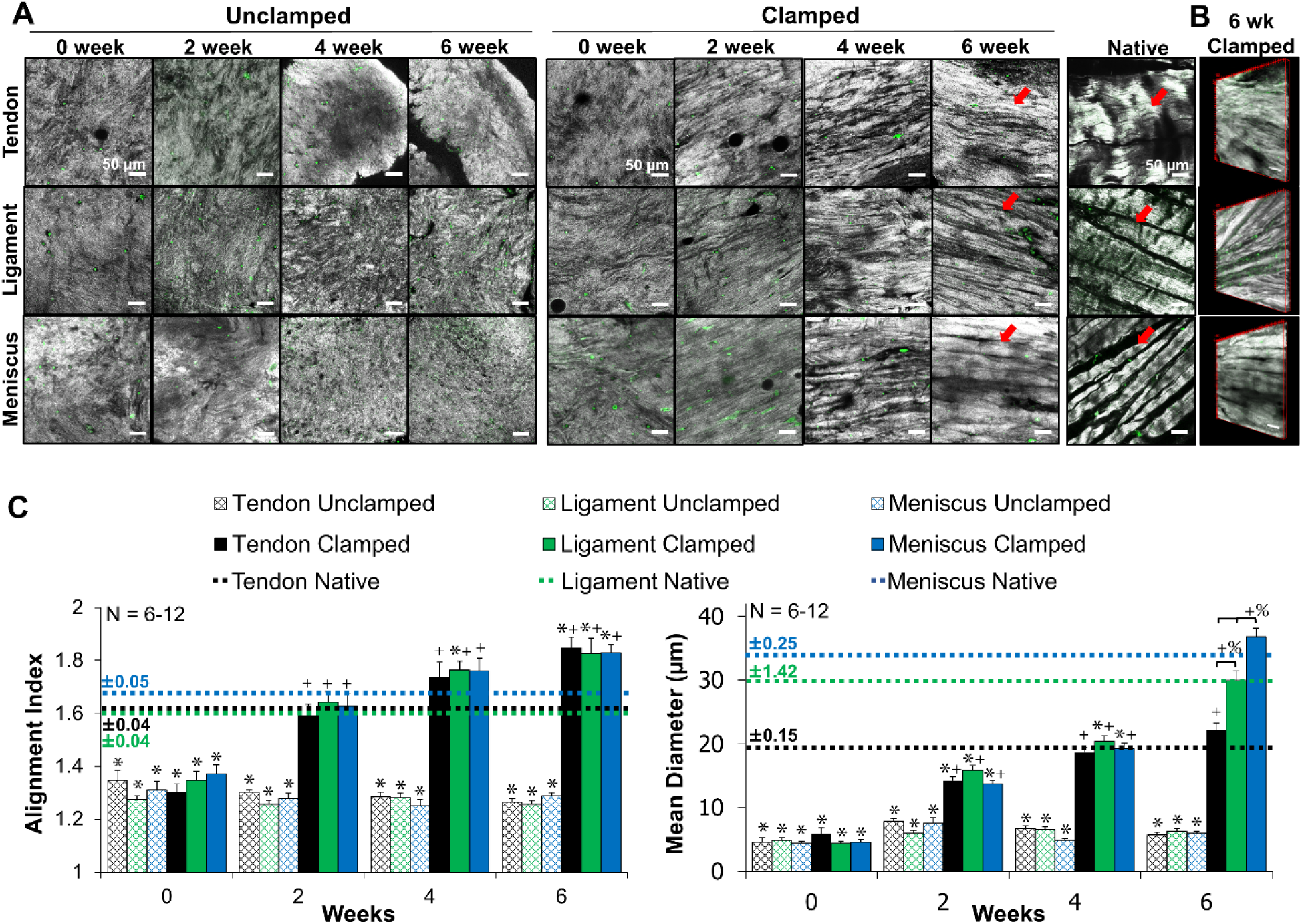
Confocal reflectance of A) collagen fiber development, B) 3D reconstruction of 6 week clamped tissues. Grey = collagen, green = auto-fluorescence of cells, scale bar = 50 µm. C) FFT based image analysis [20] of alignment (1 unorganized, 4.5 completely aligned) and mean collagen fiber diameter. Unclamped samples remained unorganized while clamped samples develop aligned fibrils by 2 weeks, which grew to significantly distinct collagen fibers (arrows) that matched respective native tissue diameters by 6 weeks. 3 to 16 images per construct were averaged, with 6-12 constructs analyzed per time point. Data shown as mean ± S.E.M., significance compared to *native, ^+^0 week, ^%^bracket group (*p* < 0.05).

Confocal reflectance performed at lower magnification (double length scale) revealed the development of hierarchical collagen organization in clamped samples, with fiber bundles or fascicles 50-350 µm in diameter (**Figure 4**). Mirroring fiber level organization, clamped samples organized similar to their respective native tissues. Specifically, tenocytes developed up to ∼50 µm diameter bundles, ligament fibroblasts develop up to ∼200 µm diameter bundles, and meniscal fibrochondrocytes developed up to ∼350 µm diameter bundles, similar to their respective tissues. Further, confocal and polarized picrosirius red imaging revealed clamped constructs began to develop crimp-like morphologies by 6 weeks, with a level of crimp that appeared to be highest in tenocyte-seeded constructs, followed by ligament fibroblasts, and finally meniscal fibrochondrocytes (**Figure 4, arrows**).

**Figure 4:**
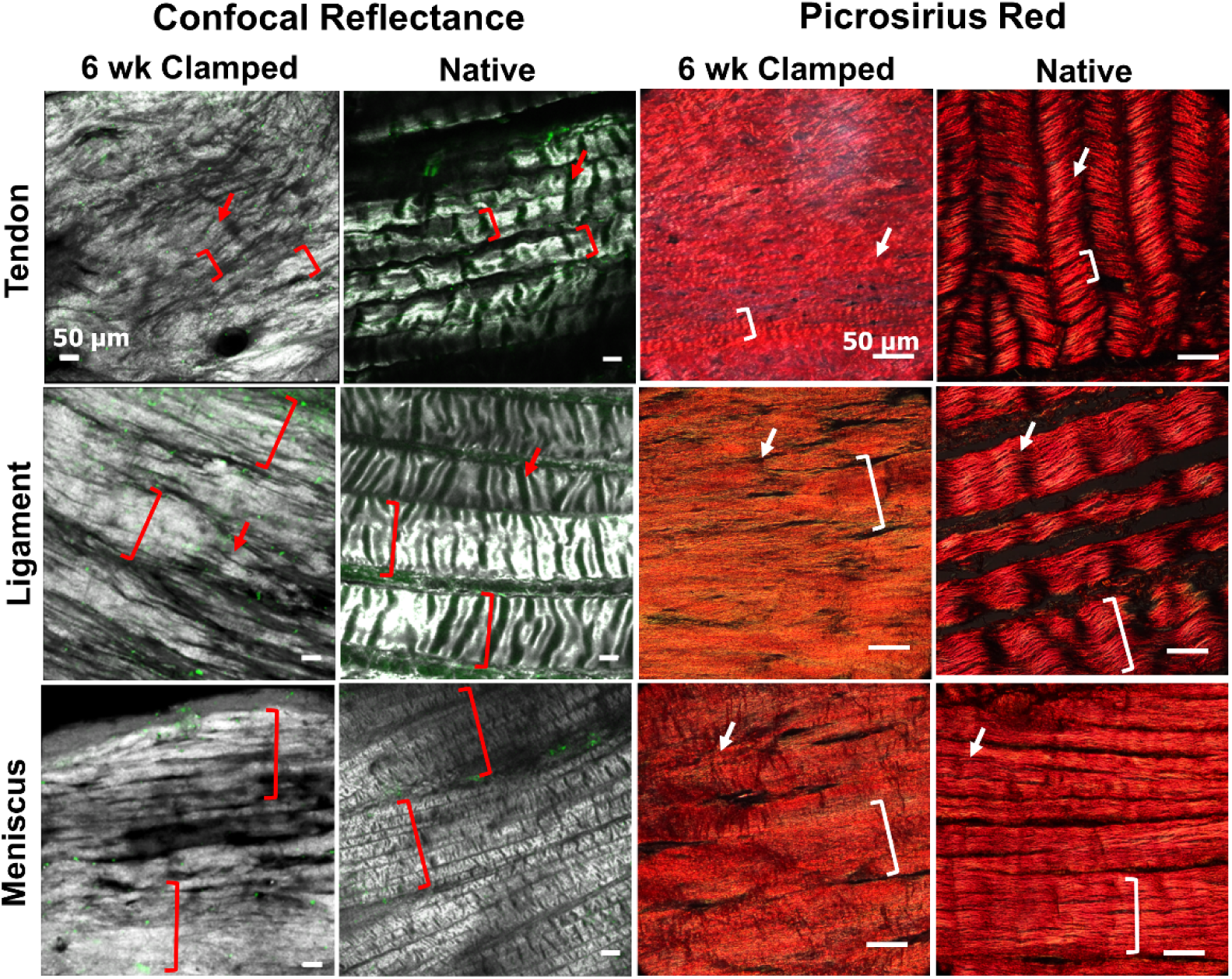
Low magnification confocal reflectance 3D reconstructions and picrosirius red stained sections imaged with polarized light, indicate that engineered samples develop hierarchical organization with fiber bundles 50-350 µm in diameter (brackets). Further, tenocytes develop strong crimp-like morphologies by 6 weeks, with similar morphologies appearing in ligament fibroblast and meniscal fibrochondrocyte cultures as well (arrows).

Higher resolution analysis with AFM demonstrated 6 week clamped samples developed some alignment at the nanometer level, but not to the extent of native tissue (**Figure 5A**). Six week clamped samples developed ∼50-90 nm wide fibrils, reaching 40-65% of native tissue diameters (**Supplemental Table 1**). Additionally, clamped samples developed ordered collagen banding, with d-period lengths ∼50-65 nm long, similar to native tissue by 6 weeks (**Figure 5B & Supplemental Table 1**), however the banding pattern in the 6 week clamped samples was less regular and well-defined compared to the native tissue samples.

**Figure 5:**
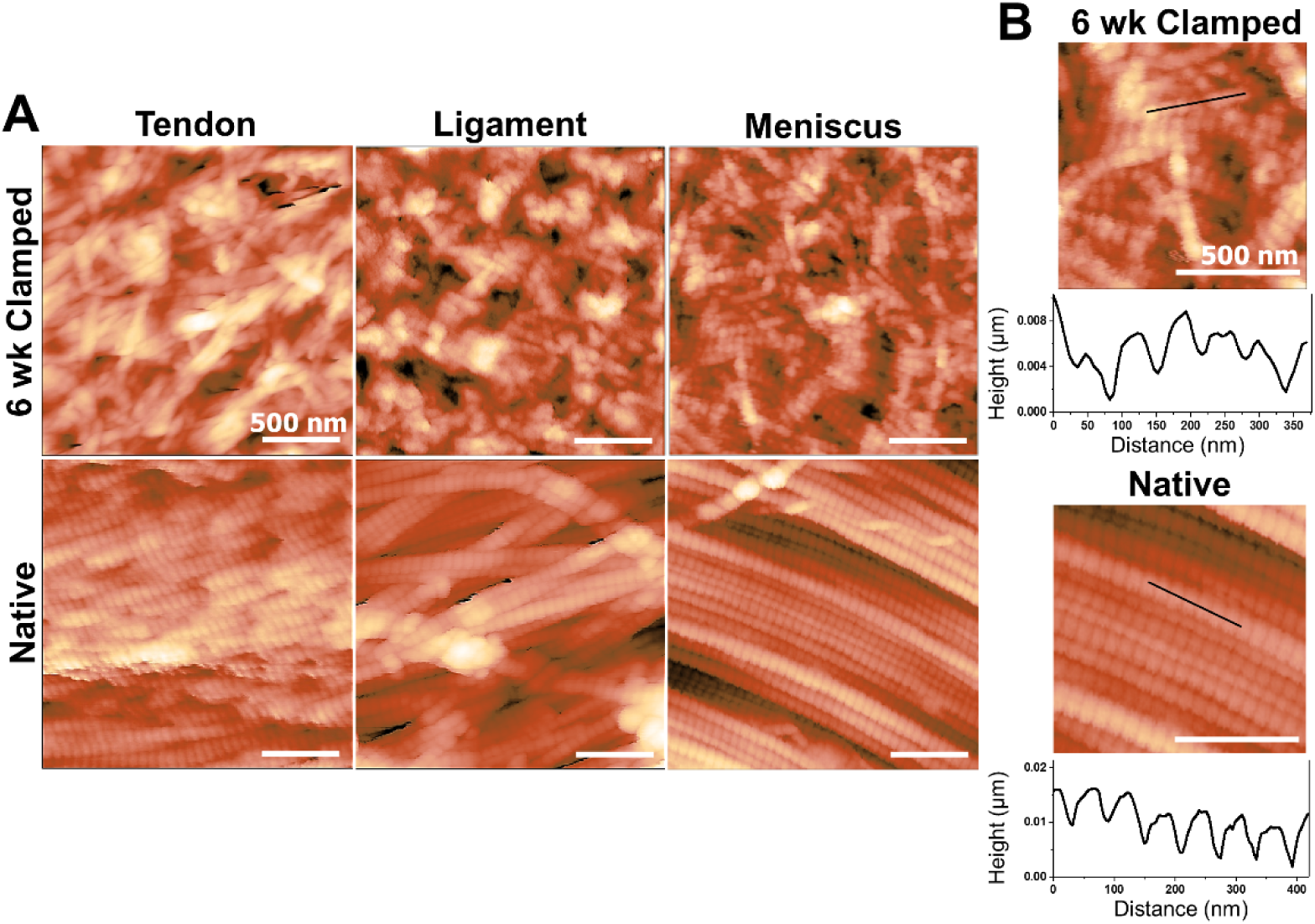
Atomic force microscopy analysis of 6 week clamped constructs and native tissue. A) Representative 2 µm x 2 µm images, B) topographical scans to determine d-period lengths. Engineered tissue developed average d-period lengths similar to native tissue by 6 weeks.

### 3.3 Tissue Biochemical Composition

Tissue level analysis revealed that DNA and collagen content were largely unchanged throughout culture. DNA per construct remained steady the entire culture for both clamped and unclamped samples, with clamped samples maintaining DNA concentrations (normalized to wet weight and dry weight) that matched respective native tissue (**Supplemental Figure 1**). Collagen content, represented by hydroxyproline (hypro), was largely unchanged throughout culture, with the exception of a significant decrease in all unclamped samples at 4 and 6 weeks (**Figure 6A**). This significant decrease in collagen content mirrors the significant contraction of unclamped samples. All clamped constructs maintained collagen concentration normalized to dry weight at 40-70% of respective native tissue throughout culture (6 week data shown, **Figure 6B**).

**Figure 6:**
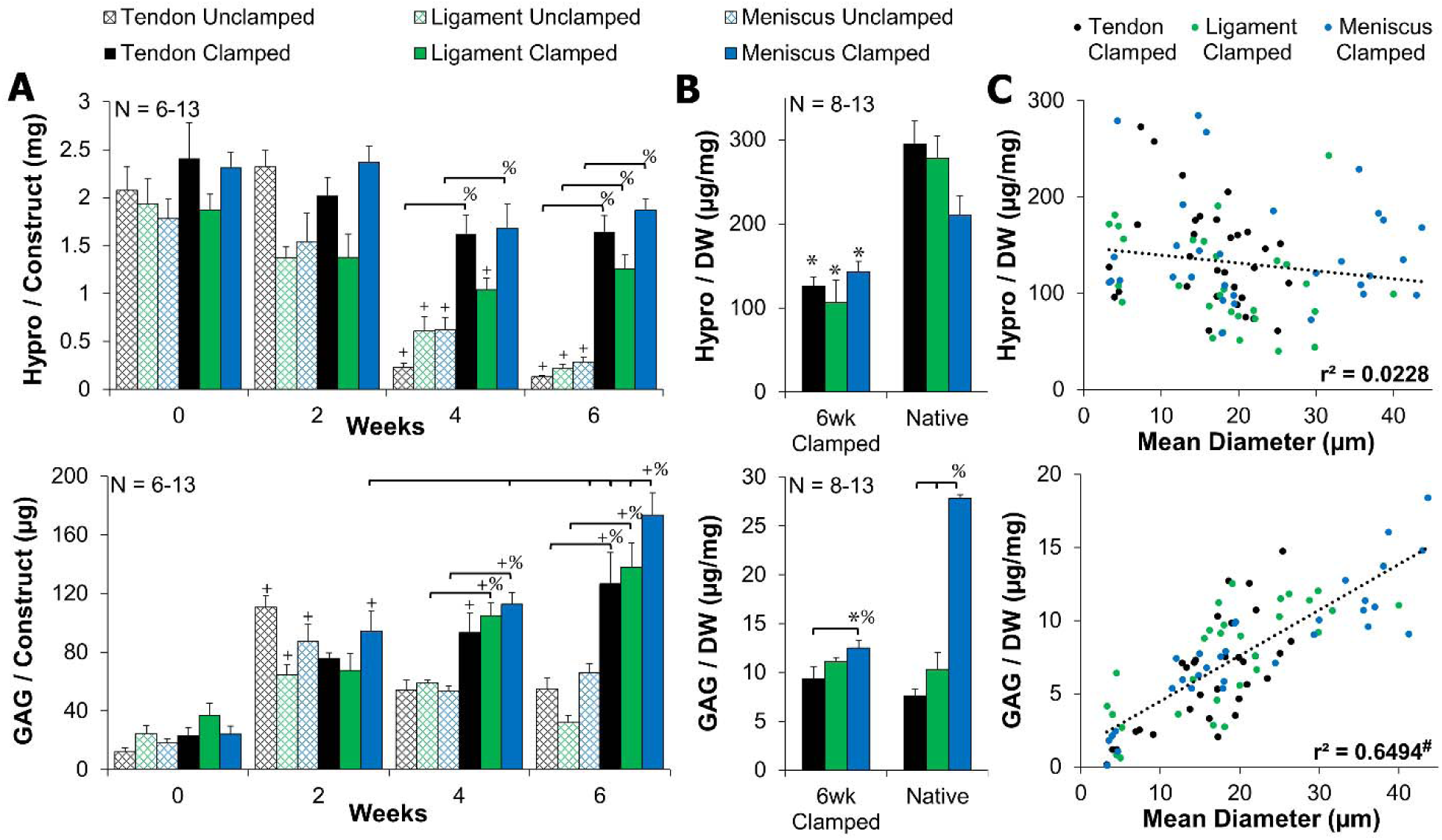
A) Clamped collagen (hydroxyproline) content was largely unchanged, while clamped GAG content significantly increased with time. B) 6 week clamped samples had 40-70% collagen/dry weight (DW) and 50-100% GAG/DW of native tissue. Data shown as mean ± S.E.M. C) Clamped GAG/DW significantly correlated with mean collagen fiber diameter, while clamped collagen/DW did not correlate. Significance compared to ^+^0 week, ^%^bracket group,*native, and ^#^significance determined by Pearson’s correlation (*p* < 0.05).

GAG content remained relatively steady in unclamped samples throughout culture. However, all clamped constructs (tendon, ligament, and meniscus) significantly increased GAG content throughout culture, with meniscal constructs having significantly more GAG accumulation than all other groups by 6 weeks (**Figure 6A**). When normalized to dry weight, tendon and ligament 6 week clamped constructs matched respective native tissue GAG concentrations (**Figure 6B**). Meniscus clamped constructs mirrored native tissue with significantly higher GAG concentrations than tendon, however it only reached 40-50% GAG content of native tissue (**Figure 6B**).

Collagen fibrillogenesis and collagen fiber size have been reported to be largely dependent on GAGs, particularly small leucine rich proteoglycans [2,28]. In this study, GAG accumulation in clamped samples appeared to mirror collagen fiber diameters suggesting GAGs are playing a role in regulating collagen fiber size. To better examine the relationship between collagen and GAG concentration on collagen fiber size, correlation analysis was performed on all clamped data from 0 to 6 weeks as cells aligned fibrils and grew them into larger µm diameter fibers. This analysis revealed a significant positive correlation between GAG concentration and mean collagen fiber size (r^2^ = 0.6494, *p* < 0.00001) and no correlation between collagen concentration and fiber size (r^2^ = 0.0228, **Figure 6C**). Examining 6 week clamped data alone, when significantly different collagen fiber sizes were produced between tendon, ligament, and meniscus constructs, this relationship was maintained, with a significant correlation between GAG content and collagen fiber size (r^2^ = 0.3235, *p* < 0.0016), and no correlation between collagen and fiber size (r^2^ = 0.0379, **Supplemental Figure 2**).

### 3.4 Tissue Mechanical Properties

Mirroring collagen fiber formation, all clamped constructs (tendon, ligament, and meniscus) significantly improved tensile moduli 20-30 fold, reaching ∼1 MPa by 6 weeks (*p* < 0.001, **Figure 7**). By 6 weeks, clamped construct tensile moduli matched reported neonatal tendon (1-2 MPa, hatched chick tendon [29]), immature ACL (1-3 MPa, 1 week old bovine [30]) and approached reported immature meniscus properties (13-26 MPa, 0-1 week old bovine [20,30]). Further, by 6 weeks all clamped samples had similar yield strength, ultimate tensile strength (UTS) and strain response (**Figure 7**). Clamped samples developed a characteristic toe region that transitioned into a linear region by ∼14-20% strain, similar to the non-linear response of tendon, ligament, and meniscus tissue. Although not significant, 6 week clamped tendon constructs trended (*p* < 0.1) to have larger strains in the toe region and at yielding compared to ligament 6 week clamped constructs, possibly due to the higher degree of crimping observed in picrosirius red and confocal analysis (**Figure 4**).

**Figure 7:**
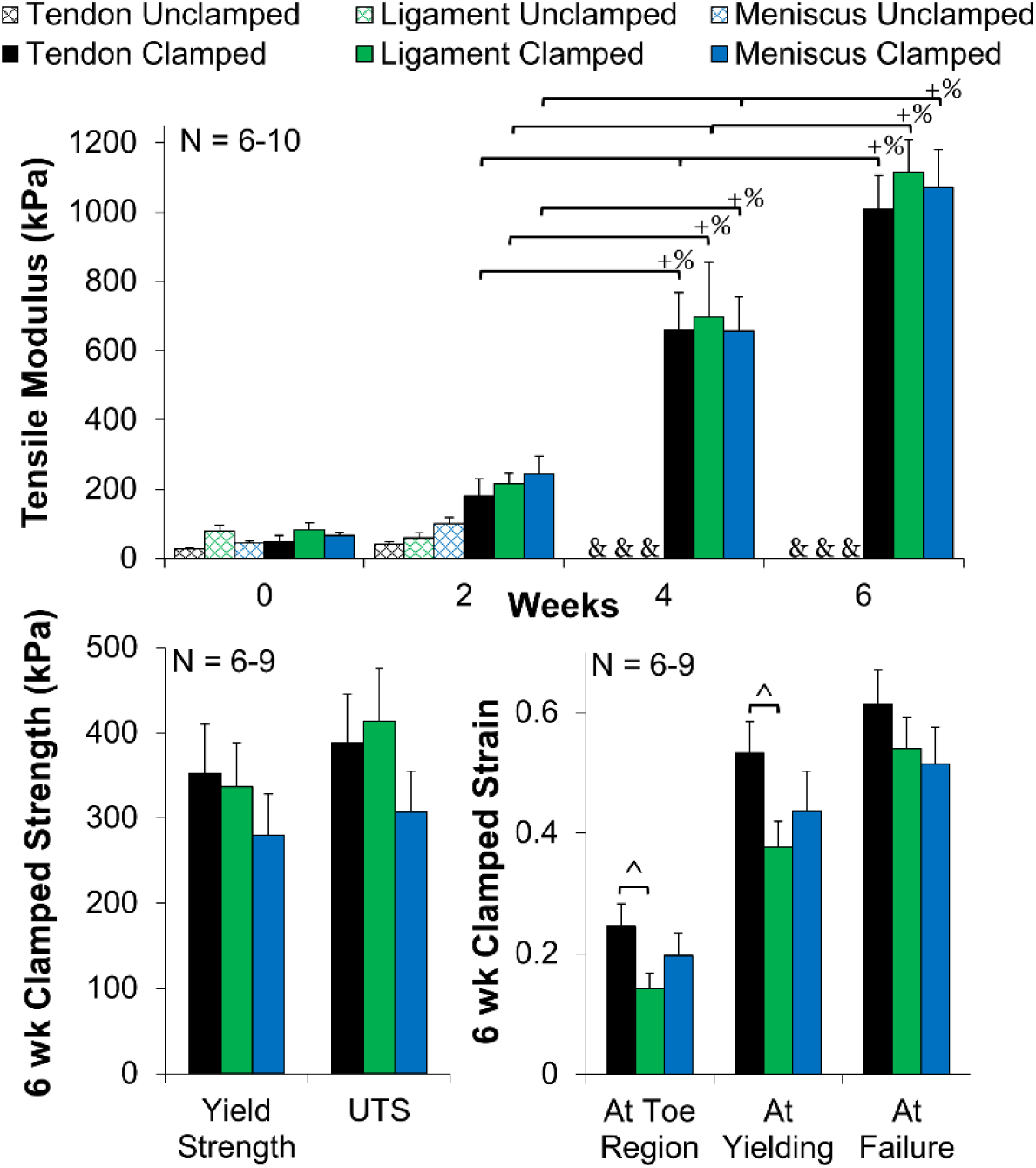
Clamped samples improved tensile moduli to ∼1 MPa by 6 week and all 6 week clamped samples had similar yield strength, ultimate tensile strength (UTS), and strain response when tested to failure at 0.75% strain/s. Tendon 6 week clamped constructs trended toward higher strains for the toe region and yielding compared to ligament. Data shown as mean ± S.E.M., significance compared to ^+^0 week, ^%^bracket group (*p* < 0.05), & denotes samples too small for mechanical analysis, ^ denotes trending (*p* < 0.1).

## 4. Discussion

This study demonstrates boundary conditions applied to cell-seeded high density collagen gels are capable of driving development of native sized and aligned hierarchical collagen fibers, 20-40 µm in diameter, with tenocytes, ligament fibroblasts, and meniscal fibrochondrocytes. The collagen fibers organize similar to native collagen from the fibril to fiber level, with native fibril banding patterns and hierarchical fiber bundles 50-350 µm in diameter by 6 weeks. Mirroring fiber organization, tensile properties of clamped samples improved significantly with time, reaching ∼1 MPa and matching native immature properties [29,30]. Additionally, cell phenotypes were maintained, with tendon, ligament, and meniscal cells differentially regulating collagen fiber formation, producing significantly different sized collagen fibers, distinct degrees of crimp, and variable GAG concentrations by 6 weeks, which all correlated to respective native tissue. To our knowledge, these are some of the largest, most hierarchically organized fibers produced to date *in vitro*.

It has been well established that mechanical boundary conditions applied to cell-seeded low density (1-5 mg/ml) collagen and fibrin gels can guide the development of embryonic-like collagen fibrils (<1 µm in diameter) [9–19]. However these systems often result in significant contraction, early rupture, low mechanical properties, and ultimately have not been shown to support development of larger fibers and fascicles *in vitro*. It has been suggested that the low concentration of collagen is a key factor in this lack of maturation [15]. In the present study, 20 mg/ml collagen gels were used, 5-20 fold higher concentration than that previously employed in low density restrained cultures. At this higher concentration, collagen fibrils (∼1 µm in diameter) align by 2 weeks of culture, similar to that previously achieved in low density systems [18], and matching native fibril tissue alignment (Figure 3). With further time in culture, fibrils mature to larger native sized fibers (20-40 µm in diameter) and fiber bundles (50-350 µm in diameter). Moreover, contraction of clamped constructs appears to reach a plateau at 4 weeks, as demonstrated by maintained size and weight between 4 and 6 weeks (Figure 2), suggesting that clamped constructs have transitioned from an alignment phase into a matrix maturation phase.

The fiber maturation in this study mirrors native fibrillogenesis in tendon, ligament, and meniscus, where collagen fibrils are believed to first align and grow longitudinally, then expand laterally [2,3]. It has been suggested that the early stages of development are intrinsically regulated, however later stages of collagen fiber maturation require mechanical stimulation [3,31,32]. As collagen concentration increases from 3 mg/ml to 20 mg/ml the degree of cellular contraction decreases and the equilibrium modulus increases significantly from 30 Pa to 1800 Pa [24]. These attributes of high density collagen gels may provide the time and mechanical signals needed to support the early maturation of collagen fibers observed in this study.

Mirroring collagen fiber organization, tensile properties improved 20-30 fold in clamped samples. The tensile modulus ultimately reached ∼1 MPa, 8-10 times higher than previous whole meniscus high-density collagen gel studies [20–22], matching neonatal tendon (1-2 MPa, hatched chick tendon [29]), immature ACL (1-3 MPa, 1 week old bovine [30]) and approaching reported immature meniscus properties (13-26 MPa, 0-1 week old bovine [20,30]). This increase, compared to the minimal change in collagen content, supports the theory that organization is more important to construct maturation than matrix accumulation [20,21,33], and stresses the importance of developing well organized collagen in engineered tissues.

Further, by employing high density collagen gels, clamped constructs maintained a collagen concentration 40-70% that of native tissue normalized to dry weight and 15-20% normalized to wet weight. In an effort to engineer tissues with higher concentrations of collagen, plastic compression of collagen gels has been investigated. In these systems, low density collagen gels are compressed, forcing water out and producing densified gels [34–37]. While these culture systems result in collagen densities similar to native tissue (100-250 mg/ml) and develop larger fiber-like morphologies, they lack the fibril level organization and do not replicate native tissue hierarchy [34–37]. In this study, the collagen concentration is 5-10 times less than that achieved in plastic compressed systems, however the tensile modulus of clamped constructs by 6 weeks matches that previously reported in plastic compression system (1-3 MPa), further stressing the importance of collagen organization in producing functional engineered tissues [36,37].

Interestingly, tendon, ligament, and meniscal constructs developed significantly different sized collagen fibers with different degrees of crimping by 6 weeks, which mirrored their native tissues. This was achieved with all the cells being cultured in the exact same environment, suggesting cells intrinsically regulate fiber development. It is well established that fibril organization is intrinsically regulated; however higher level fiber and fascicle organizations are thought to be more influenced by the mechanical environment [3,31,32]. Here we demonstrate that meniscal fibrochondrocytes, ACL ligament fibroblasts, and flexor tenocytes will form tissue specific fiber and fascicle organization when cultured with the same static load boundary conditions. Understanding these cellular differences is key to successfully driving fiber development in engineered replacement and *in vivo* after repair.

Cells regulate fibril and fiber formation by many means, including cytoskeletal tension, integrins, fibronectin, and multiple macromolecules, including small leucine-rich proteoglycans (SLRPs) and multiple types of collagen [2,18,28]. Tenocytes and ligament fibroblasts have been shown to have a tensional homeostasis regulated by α-smooth muscle actin, at which gene expression changes from catabolic to anabolic [11,38]. In this study, all clamped samples had significant contraction between ∼2-4 weeks, which stabilized by 4 weeks. Further, the largest growth in collagen fiber size was observed between 4-6 weeks, suggesting cells have reached a tensional homeostasis at 4 weeks and transitioned to cell specific anabolic signaling to drive fiber formation.

Intriguingly, in this study GAG/proteoglycan accumulation mirrors fiber diameters, with a significant positive correlation between the two properties (r^2^ = 0.65), suggesting GAGs are playing a role in fiber regulation. The major proteoglycans of tendon and ligaments are SLRPs, including decorin, biglycan, and fibromodulin, which are known to play a large, complex role in collagen fiber size [2,3,28]. By 6 weeks, tendon and ligament constructs matched respective native tissue GAG concentrations and fiber sizes, suggesting accumulation of SLRPs. Meniscus constructs only reach ∼50% GAG accumulation of native tissue, however menisci have significantly more GAG than tendons and ligaments, composed primarily of large aggrecan molecules. Aggrecan accumulation is relatively low during development and localized to the inner 1/3 of the tissue where large hierarchical collagen fibers do not form [39], possibly accounting for the lack of GAG in engineered tissues compared to native tissue in this study. Nonetheless, the positive correlation between fiber size and GAG accumulation may be skewed by this tendency of meniscal fibrochondrocytes to create more aggrecan and not necessarily SLRPs such as decorin, biglycan, and fibromodulin. The positive correlation between GAGs and fiber size in this study only begins to scratch the surface of how cells regulate their fiber sizes and future studies should more closely evaluate specific SLRP accumulation and localization.

In addition to significantly different fiber sizes, tendon, ligament, and meniscal constructs developed varying degrees of crimp. Tendon constructs appear to develop the largest degree of crimp by 6 weeks, mirroring native tissue variances. Crimps are wave-like morphologies in collagen fibrils and fibers that play an important role in force transduction by forming the toe region of stress-strain curve [40]. In this study, tendon constructs, which exhibited the largest amount of crimp (Figure 4), trended toward having larger toe regions than ligament constructs (Figure 7), suggesting the crimp is playing a functional role and providing a more native-like non-linear mechanical response.

Crimp is known to vary in degree and angle between tendons and ligaments throughout the body. It first appears during embryonic development [40] and it increases in wavelength with increasing strain and maturation [40,41]. Crimp has been shown to be generated by cellular contractile forces [42,43] and regulated by differences in stiffness between fibrils and matrix [42]. While it is still largely debated how crimps form, they are believed to be a clear reflection of the local mechanobiological environment of the extracellular matrix [43].

To our knowledge, crimp has not been observed in low density constructs while they are constrained, suggesting these systems do not reach a mechanical threshold necessary for cells to imitate crimp formation. In this study, we observe crimp formation by 6 weeks, suggesting high density collagen gels support fiber maturation to the mechanical threshold necessary for crimp formation. Further, differences in degree of crimp between cell types demonstrate that cells have different levels of contraction and mechanical sensing. Collectively, differences in fiber size, GAG accumulation, and crimp observed in this study suggest differences in cellular contractility and mechanical sensing. This system provides a platform to further evaluate how cellular mechanical sensing and integrins regulates fiber and fascicle maturation. Understanding this mechanobiological relationship may help to drive repair and regeneration after injury and maintain homeostasis of fibrous musculoskeletal tissues throughout the body. Future studies should investigate differences in actin expression and changes in gene expression with time in culture to better explore the cell-matrix interactions during collagen fiber development.

Collectively this study demonstrates the potential of high density collagen gels, cultured with mechanical boundary conditions, to drive hierarchical collagen fiber formation. Despite the significant improvements in fiber organization and mechanical properties observed in this study, there is still further fiber maturation needed to match native tissue properties. On the nanometer scale, native fibril alignment was lacking and fibrils reached only ∼40-70% diameter of native tissues. Additionally, while collagen bundles, 50-350 µm in diameter, resembled early fascicle development, evaluation of the next hierarchical level would reveal a need for further maturation. Similarly, while tensile moduli match native immature tissue values, further improvements are needed to match adult tissue modulus, ultimate tensile strength, and strain properties. Collectively, this suggests the need for further development of collagen fibers. This system sets a solid base for integration of additional mechanical [4,6,7,21,44] or chemical signals [4,6,7,45,46] to improve density, crosslinking, and maturation of fascicles. Despite the need for further maturation to serve as functional replacements, cells produced tissue specific collagen organization and GAG accumulation, suggesting this system could serve as a viable platform for investigating cellular regulation of collagen fiber organization, diseases, and injury *in vitro*.

## 5. Conclusions

This study provides new insight into how cell develop collagen fibers *in vitro* and provides a method for developing some of the most organized collagen scaffolds to date for tendon, ligament, and meniscus tissue. This model could be a promising tool to investigate collagen fiber development, disease, and injury *in vitro*, and these constructs demonstrate great promise as functional musculoskeletal replacements.

## Author Contributions

J.L.P conceived and designed the experiments. J.L.P, T.M, and I.S. performed experiments and analyzed the data. A.G. performed AFM analysis. M.M.S. supervised the project. J.L.P., T.M., I.S., A.G., and M.M.S. wrote and edited manuscript.

## Data Availability

Research data are available upon request from

## Acknowledgements

J.L.P. was supported by the Whitaker International Program, Institute of International Education, USA. J.L.P. and M.M.S. where funded by the grant from the UK Regenerative Medicine Platform “ Acellular Approaches for Therapeutic Delivery” (MR/K026682/1). A.G. acknowledges support from the European Union’s Horizon 2020 Research and Innovation Programme through the Marie Skłodowska-Curie Individual Fellowship “ RAISED” [660757]. A.G. and M.M.S. acknowledge support from a Wellcome Trust Senior Investigator Award (098411/Z/12/Z).

M.M.S. acknowledges the grant “ State of the Art Biomaterials Development and Characterization of the Cell-Biomaterials Development and Characterization of the Cell-Biomaterial Interface” (MR/L012677/1) from the MRC, the ERC Seventh Framework Programme Consolidator grant “ Natural CG’ [616417], and the grant from the UK Regenerative Medicine Platform “ Acellular/Smart Materials – 3D Architecture” [MR/R015651/1]. The authors acknowledge the use of the Facility for Imaging and Light Microscopy (FILM) at Imperial College London.

## References

[1] M.J. Buehler, Nature designs tough collagen: Explaining the nanostructure of collagen fibrils, Proc. Natl. Acad. Sci. (2006). https://doi.org/10.1073/pnas.0603216103.

[2] G. Zhang, B.B. Young, Y. Ezura, M. Favata, L.J. Soslowsky, S. Chakravarti, D.E. Birk, Development of tendon structure and function: regulation of collagen fibrillogenesis., J. Musculoskelet. Neuronal Interact. 5 (2005) 5–21. http://www.ncbi.nlm.nih.gov/pubmed/15788867.

[3] B.K. Connizzo, S.M. Yannascoli, L.J. Soslowsky, Structure-function relationships of postnatal tendon development: A parallel to healing, Matrix Biol. 32 (2013) 106–116. https://doi.org/10.1016/j.matbio.2013.01.007.

[4] M.T. Rodrigues, R.L. Reis, M.E. Gomes, Engineering tendon and ligament tissues: Present developments towards successful clinical products, J. Tissue Eng. Regen. Med. (2013). https://doi.org/10.1002/term.1459.

[5] W. Petersen, B. Tillmann, Collagenous fibril texture of the human knee joint menisci, Anat. Embryol. (Berl). 197 (1998) 317–324. https://doi.org/10.1007/s004290050141.

[6] J. Hasan, J. Fisher, E. Ingham, Current strategies in meniscal regeneration, J. Biomed. Mater. Res. - Part B Appl. Biomater. 102 (2014) 619–634. https://doi.org/10.1002/jbm.b.33030.

[7] T. Nau, A. Teuschl, Regeneration of the anterior cruciate ligament: Current strategies in tissue engineering, World J. Orthop. 6 (2015) 127. https://doi.org/10.5312/wjo.v6.i1.127.

[8] S. Patel, J.M. Caldwell, S.B. Doty, W.N. Levine, S. Rodeo, L.J. Soslowsky, S. Thomopoulos, H.H. Lu, Integrating soft and hard tissues via interface tissue engineering, J. Orthop. Res. 36 (2018) 1069–1077. https://doi.org/10.1002/jor.23810.

[9] E. Bell, B. Ivarsson, C. Merrill, Production of a tissue-like structure by contraction of collagen lattices by human fibroblasts of different proliferative potential in vitro., Proc. Natl. Acad. Sci. U. S. A. 76 (1979) 1274–8. http://www.ncbi.nlm.nih.gov/pubmed/286310%0Ahttp://www.pubmedcentral.nih.gov/articlerender.fcgi?artid=PMC383233.

[10] K.D. Costa, E.J. Lee, J.W. Holmes, Creating Alignment and Anisotropy in Engineered Heart Tissue: Role of Boundary Conditions in a Model Three-Dimensional Culture System, Tissue Eng. 9 (2003) 567–577. https://doi.org/10.1089/107632703768247278.

[11] D.R. Henshaw, E. Attia, M. Bhargava, J.A. Hannafin, Canine ACL fibroblast integrin expression and cell alignment in response to cyclic tensile strain in three-dimensional collagen gels, J. Orthop. Res. (2006). https://doi.org/10.1002/jor.20050.

[12] R.D. Bowles, R.M. Williams, W.R. Zipfel, L.J. Bonassar, Self-Assembly of Aligned Tissue-Engineered Annulus Fibrosus and Intervertebral Disc Composite Via Collagen Gel Contraction, Tissue Eng. Part A. 16 (2010) 1339–1348.

[13] S. Thomopoulos, G.M. Fomovsky, J.W. Holmes, The Development of Structural and Mechanical Anisotropy in Fibroblast Populated Collagen Gels, J. Biomech. Eng. 127 (2005) 742. https://doi.org/10.1115/1.1992525.

[14] H.A. Awad, D.L. Butler, M.T. Harris, R.E. Ibrahim, Y. Wu, R.G. Young, S. Kadiyala, G.P. Boivin, In vitro characterization of mesenchymal stem cell-seeded collagen scaffolds for tendon repair: Effects of initial seeding density on contraction kinetics, J. Biomed. Mater. Res. (2000). https://doi.org/10.1002/(SICI)1097-4636(200008)51:2<233::AID-JBM12>3.0.CO;2-B.

[15] A. Herchenhan, M.L. Bayer, R.B. Svensson, S.P. Magnusson, M. Kjær, In vitro tendon tissue development from human fibroblasts demonstrates collagen fibril diameter growth associated with a rise in mechanical strength, Dev. Dyn. 242 (2013) 2–8. https://doi.org/10.1002/dvdy.23896.

[16] A.P. Breidenbach, N.A. Dyment, Y. Lu, M. Rao, J.T. Shearn, D.W. Rowe, K.E. Kadler, D.L. Butler, Fibrin Gels Exhibit Improved Biological, Structural, and Mechanical Properties Compared with Collagen Gels in Cell-Based Tendon Tissue-Engineered Constructs, Tissue Eng. Part A. (2014). https://doi.org/10.1089/ten.tea.2013.0768.

[17] V.S. Nirmalanandhan, M. Rao, M.S. Sacks, B. Haridas, D.L. Butler, Effect of length of the engineered tendon construct on its structure-function relationships in culture, J. Biomech. (2007). https://doi.org/10.1016/j.jbiomech.2006.11.016.

[18] M.L. Bayer, C.Y.C. Yeung, K.E. Kadler, K. Qvortrup, K. Baar, R.B. Svensson, S. Peter Magnusson, M. Krogsgaard, M. Koch, M. Kjaer, The initiation of embryonic-like collagen fibrillogenesis by adult human tendon fibroblasts when cultured under tension, Biomaterials. 31 (2010) 4889–4897. https://doi.org/10.1016/j.biomaterials.2010.02.062.

[19] J.Z. Paxton, L.M. Grover, K. Baar, Engineering an In Vitro Model of a Functional Ligament from Bone to Bone, Tissue Eng. Part A. 16 (2010) 3515–3525. https://doi.org/10.1089/ten.tea.2010.0039.

[20] J.L. Puetzer, E. Koo, L.J. Bonassar, Induction of fiber alignment and mechanical anisotropy in tissue engineered menisci with mechanical anchoring, J. Biomech. 48 (2015) 1967. https://doi.org/10.1016/j.jbiomech.2015.02.033.

[21] J. Puetzer, L. Bonassar, Physiologically Distributed Loading Patterns Drive the Formation of Zonally Organized Collagen Structures in Tissue-Engineered Meniscus, Tissue Eng. Part A. 22 (2016) 907–916. https://doi.org/10.1089/ten.tea.2015.0519.

[22] M.C. McCorry, L.J. Bonassar, Fiber Development and Matrix Production in Tissue Engineered Menisci using Bovine Mesenchymal Stem Cells and Fibrochondrocytes, Connect. Tissue Res. 58 (2017) 329–341. https://doi.org/10.1002/cncr.27633.Percutaneous.

[23] J.L. Puetzer, L.J. Bonassar, High density type I collagen gels for tissue engineering of whole menisci, Acta Biomater. 9 (2013) 7787–7795. https://doi.org/10.1016/j.actbio.2013.05.002.

[24] V.L. Cross, Y. Zheng, N. Won Choi, S.S. Verbridge, B.A. Sutermaster, L.J. Bonassar, C. Fischbach, A.D. Stroock, Dense type I collagen matrices that support cellular remodeling and microfabrication for studies of tumor angiogenesis and vasculogenesis in vitro, Biomaterials. 31 (2010) 8596–8607. https://doi.org/10.1016/j.biomaterials.2010.07.072.

[25] M.S. Bergholt, J.-P. St-Pierre, G.S. Offeddu, P.A. Parmar, M.B. Albro, J.L. Puetzer, M.L. Oyen, M.M. Stevens, Raman spectroscopy reveals new insights into the zonal organization of native and tissue-engineered articular cartilage, ACS Cent. Sci. 2 (2016). https://doi.org/10.1021/acscentsci.6b00222.

[26] R. Neuman, M. Logan, The determination of Hydroxyproline, J. Biol. Chem. 184 (1950) 299–306. http://www.ncbi.nlm.nih.gov/pubmed/15421999.

[27] B.O. Enobakhare, D.L. Bader, D.A. Lee, Quantification of Sulfated Glycosaminoglycans in Chondrocyte/Alginate Cultures, by Use of 1,9-Dimethylmethylene Blue, Anal. Biochem. 243 (1996) 189–191. https://doi.org/10.1006/abio.1996.0502.

[28] S. Chen, D.E. Birk, The regulatory roles of small leucine-rich proteoglycans in extracellular matrix assembly, FEBS J. 280 (2013) 2120–2137. https://doi.org/10.1111/febs.12136.

[29] D.J. McBride, R.L. Trelstad, F.H. Silver, Structural and mechanical assessment of developing chick tendon, Int. J. Biol. Macromol. (1988). https://doi.org/10.1016/0141-8130(88)90048-7.

[30] S. V. Eleswarapu, D.J. Responte, K.A. Athanasiou, Tensile properties, collagen content, and crosslinks in connective tissues of the immature knee joint, PLoS One. 6 (2011) 1–7. https://doi.org/10.1371/journal.pone.0026178.

[31] B. Mikic, T.L. Johnson, A.B. Chhabra, B.J. Schalet, M. Wong, E.B. Hunziker, Differential effects of embryonic immobilization on the development of fibrocartilaginous, J. Rehabil. Res. Dev. 37 (2000) 127–133.

[32] A.G. Schwartz, J.H. Lipner, J.D. Pasteris, G.M. Genin, S. Thomopoulos, Muscle loading is necessary for the formation of a functional tendon enthesis, Bone. 55 (2013) 44–51. https://doi.org/10.1016/j.bone.2013.03.010.

[33] B.M. Baker, R.L. Mauck, The effect of nanofiber alignment on the maturation of engineered meniscus constructs., Biomaterials. 28 (2007) 1967–77. https://doi.org/10.1016/j.biomaterials.2007.01.004.

[34] Y. Wang, J. Silvent, M. Robin, F. Babonneau, A. Meddahi-Pellé, N. Nassif, M.M. Giraud Guille, Controlled collagen assembly to build dense tissue-like materials for tissue engineering, Soft Matter. 7 (2011) 9659–9664. https://doi.org/10.1039/c1sm05868a.

[35] T. Novak, B. Seelbinder, C.M. Twitchell, C.C. Van Donkelaar, S.L. Voytik-Harbin, C.P. Neu, Mechanisms and Microenvironment Investigation of Cellularized High Density Gradient Collagen Matrices via Densification, Adv. Funct. Mater. (2016). https://doi.org/10.1002/adfm.201503971.

[36] J.L. Zitnay, S.P. Reese, G. Tran, N. Farhang, R.D. Bowles, J.A. Weiss, Fabrication of dense anisotropic collagen scaffolds using biaxial compression, Acta Biomater. 65 (2018) 76–87. https://doi.org/10.1016/j.actbio.2017.11.017.

[37] R.A. Brown, M. Wiseman, C.B. Chuo, U. Cheema, S.N. Nazhat, Ultrarapid engineering of biomimetic materials and tissues: Fabrication of nano- and microstructures by plastic compression, Adv. Funct. Mater. (2005). https://doi.org/10.1002/adfm.200500042.

[38] M. Lavagnino, S.P. Arnoczky, In vitro alterations in cytoskeletal tensional homeostasis control gene expression in tendon cells, J. Orthop. Res. (2005). https://doi.org/10.1016/j.orthres.2005.04.001.

[39] J. Melrose, S. Smith, M. Cake, R. Read, J. Whitelock, Comparative spatial and temporal localisation of perlecan, aggrecan and type I, II and IV collagen in the ovine meniscus: An ageing study, Histochem. Cell Biol. (2005). https://doi.org/10.1007/s00418-005-0005-0.

[40] N.S. Kalson, Y. Lu, S.H. Taylor, D.F. Holmes, K.E. Kadler, A structure-based extracellular matrix expansion mechanism of fibrous tissue growth, Elife. (2015). https://doi.org/10.7554/eLife.05958.

[41] T.A.H. Järvinen, L. Jozsa, P. Kannus, T.L.N. Järvinen, M. Kvist, T. Hurme, J. Isola, H. Kalimo, M. Järvinen, Mechanical loading regulates tenascin-C expression in the osteotendinous junction, J. Cell Sci. (1999).

[42] A. Herchenhan, N.S. Kalson, D.F. Holmes, P. Hill, K.E. Kadler, L. Margetts, Tenocyte contraction induces crimp formation in tendon-like tissue, Biomech. Model. Mechanobiol. 11 (2012) 449–459. https://doi.org/10.1007/s10237-011-0324-0.

[43] M. Lavagnino, A.E. Brooks, A.N. Oslapas, K.L. Gardner, S.P. Arnoczky, Crimp length decreases in lax tendons due to cytoskeletal tension, but is restored with tensional homeostasis, J. Orthop. Res. (2017). https://doi.org/10.1002/jor.23489.

[44] N.S. Kalson, D.F. Holmes, A. Herchenhan, Y. Lu, T. Starborg, K.E. Kadler, Slow stretching that mimics embryonic growth rate stimulates structural and mechanical development of tendon-like tissue in vitro, Dev. Dyn. (2011). https://doi.org/10.1002/dvdy.22760.

[45] L.M. Delgado, K. Fuller, D.I. Zeugolis, Collagen Cross-Linking: Biophysical, Biochemical, and Biological Response Analysis, Tissue Eng. - Part A. (2017). https://doi.org/10.1089/ten.tea.2016.0415.

[46] E.A. Makris, D.J. Responte, N.K. Paschos, J.C. Hu, K.A. Athanasiou, Developing functional musculoskeletal tissues through hypoxia and lysyl oxidase-induced collagen cross-linking, Proc. Natl. Acad. Sci. 111 (2014) E4832–E4841. https://doi.org/10.1073/pnas.1414271111.

