## Supplemental for "Driving Hierarchical Collagen Fiber Formation for Functional Tendon, Ligament, and Meniscus Replacement"

### Supplemental Table 1

| | | D-period Length (nm) | Fibril Diameter (nm) |
|---|---|---|---|
| **Tendon** | 6 wk Clamped | 56.3 ± 6.1 | 89.3 ± 19.2* |
| | Native | 61.7 ± 1.2 | 136 ± 14.4 |
| **Ligament** | 6 wk Clamped | 59.3 ± 7.7 | 62.3 ± 15.1* |
| | Native | 66.1 ± 1.4 | 126 ± 13.3 |
| **Meniscus** | 6 wk Clamped | 60.6 ± 4.1 | 54.7 ± 13.1* |
| | Native | 61.2 ± 0.5 | 131 ± 8.8 |

**Supplemental Table 1:** Fibril measurements of d-period length (banding) and diameter from AFM analysis of 6 week clamped and native tissue. Banding lengths were similar between engineered and native tissues, while engineered tissue only reached 40-65% diameter of native tissue. Measurements were taken from 5-6 fibrils per sample and pooled for general analysis of fibril characteristics (n = 10-15 for engineered, and n = 5-6 for native). * Significance compared to respective native tissue ($p < 0.05$).

### Supplemental Figures

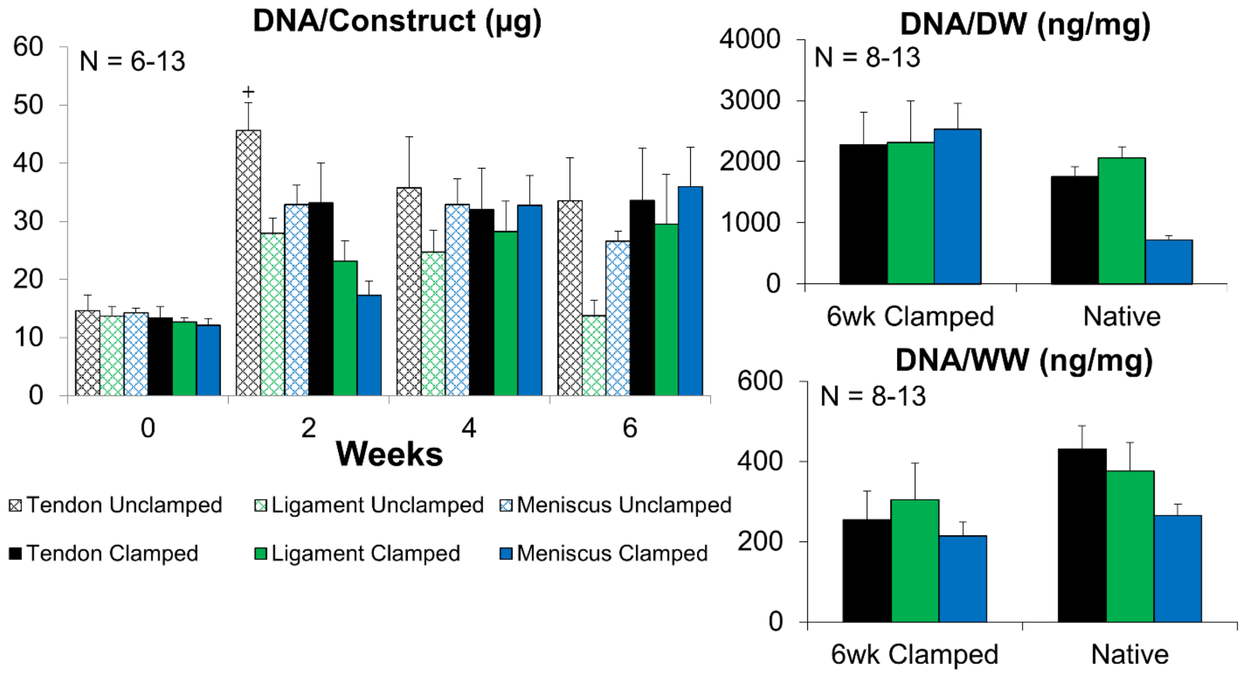


**Supplemental Figure 1:** DNA of clamped and unclamped samples remained relatively constant throughout culture, with 6 week clamped samples having no significant differences from native tissue DNA concentrations normalized to wet weight (WW) and dry weight (DW). Data shown as mean ± standard error. Significance compared to $^{+}$0 week ($p$ < 0.05).

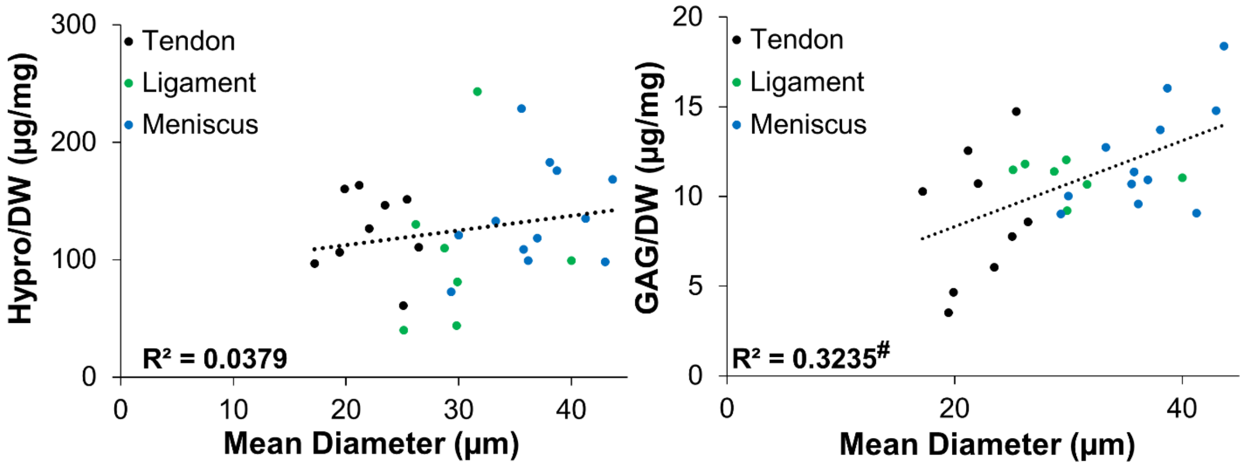


**Supplemental Figure 2:** Correlation analysis of 6 week clamped sample collagen (Hypro) and GAG content normalized to dry weight (DW) compared to mean collagen fiber diameter. Clamped 6 week construct GAG/DW significantly correlated with collagen fiber size, while Collagen (hypro) / DW did not correlate with fiber size. $^{\#}$Significance determined by Pearson's correlation.
